# YAP1-TAZ/TEAD transcriptional networks restrain differentiation downstream of oncogenic Hedgehog-SMO activity

**DOI:** 10.1101/2020.12.28.424593

**Authors:** Yao Yuan, Natalia Salinas Parra, Qianming Chen, Ramiro Iglesias-Bartolome

## Abstract

Disruption of the transcriptional activity of the Hippo pathway members YAP1 and TAZ has become a major target for cancer treatment. However, detailed analysis of the effectivity and networks affected by YAP1/TAZ transcriptional targeting are limited. Here, by comparing the effects of YAP1/TAZ knockdown with those resulting from TEAD blockage, we unveil the consequences of YAP1/TAZ transcriptional inhibition in cancer cells. We utilize TEADi, an inhibitor of the binding of YAP1 and TAZ with their main transcriptional target TEAD. In a mouse model of basal cell carcinoma (BCC) driven by the smoothened oncogene (SmoM2), TEADi and YAP1/TAZ knockdown lead to reduced proliferation and increased differentiation of tumor cells both in vitro and in vivo. We find that TEAD transcriptional networks inactivate differentiation in BCC by regulating KLF4. Furthermore, we determine YAP1/TAZ TEAD-independent effects in cancer cells that impact Stat3 and NF-κB gene networks. Our results reveal the TEAD dependent and independent roles of YAP1/TAZ in cancer and expose potential pitfalls for targeting TEAD transcription in tumors.

## Introduction

The Hippo pathway is one of the main signaling networks altered in human cancer ^1^, with genomic and signaling changes in Hippo components leading to activation and nuclear translocation of YAP1 and TAZ (*WWTR1*). YAP1 and TAZ are paralog proteins that interact with a wide range of cytoplasmic and nuclear transcriptional effectors, ultimately modulating the proliferative and differentiation status of cells in response to chemical and mechanical microenvironmental cues ^2,3^.

In the skin, YAP1 and TAZ are activated in basal (BCC) and squamous (SCC) cell carcinoma ^4^. Interestingly, while YAP1 epidermal knockout leads to a reduction in tumor lesions in a mouse model of BCC ^5^, a more profound effect is observed with the concomitant downregulation of both YAP1 and TAZ in mouse BCC and SCC tumors ^6^. This evidence indicates that disruption of both YAP1 and TAZ signaling might be necessary to achieve significant responses for BCC and SCC treatment. One common YAP1 and TAZ downstream pathway is the activation of TEAD transcription factors and efforts are underway to develop YAP1/TAZ-TEAD interaction inhibitors that could be used for cancer and other hyperproliferative diseases. However, a major challenge in studying the effectivity of this approach is the lack of preclinical models to characterize the consequences of TEAD inhibition.

Precise evidence of the effectiveness of targeting YAP1/TAZ-TEAD has been limited to the use of verteporfin ^7^ and peptide inhibitors ^8,9^. One problem with this approach is that verteporfin has YAP1/TAZ-TEAD independent effects ^10,11^, and drugs and soluble peptide inhibitors do not provide cellular and tissue level information. On the other hand, studies of the role of YAP1/TAZ in cancer involve the knockout of these proteins, affecting TEAD-dependent events and other transcriptional and signaling components that interact with YAP1 and TAZ ^12,13^. This complicates the precise understanding of TEAD functions and masks the potential differences between absence versus blockage of YAP1/TAZ proteins. An additional tool to study TEAD specificity are rescue experiments with the YAP1 S94/F95 mutant that does not bind to TEAD ^14^, although YAP1 and TAZ have to be knocked-down or knocked-out for these assays and the rescue leads to YAP1 overexpression.

To circumvent some of the limitations to study TEAD inhibition in cells and tissues our group developed TEADi, a genetically encoded fluorescently traceable dominant negative protein that blocks nuclear interaction of TEAD with YAP1 and TAZ ^15^. TEADi presents several advantages, including rapid inhibition of TEAD transcription and concomitant blockage of both YAP1 and TAZ without altering structural or cytoplasmic functions of these proteins. TEADi can be used to dissect in more detail the consequences of TEAD blockage and could serve as a resource to differentiate TEAD-dependent and independent effects, providing additional clues to suppress YAP1/TAZ activity in cancer.

Here, we utilize TEADi in a mouse model of BCC driven by oncogenic Hedgehog-Smoothened (SmoM2) activity to analyze the transcriptional and cell fate consequences of TEAD blockage in skin cancer. We find that TEAD inhibition in BCC triggers the rapid activation of differentiation programs, both in cell culture and in mouse skin. The activation of differentiation gene networks downstream of TEAD inhibition is dependent on the activation of KLF4. By comparing the effects of TEADi with those triggered by YAP1 and TAZ knockdown, our results reveal the TEAD dependent and independent roles of YAP1/TAZ in tumors and the potential issues for targeting TEAD transcription in cancer.

## Results

### YAP1/TAZ-TEAD regulate differentiation gene networks in BCC

For our studies we utilized a mouse model of BCC driven by expression of the constitutive active SmoM2 oncogene. Lox-stop-lox (LSL)-SmoM2 mice ^16^ were crossed with mice carrying a tamoxifen inducible cre-recombinase under the control of the cytokeratin 14 promoter (K14CreERT) ^17^ to target basal cells in the skin (from now on named K14-SmoM2) (Fig. 1A). As it has been demonstrated before ^18^, tamoxifen treatment in K14-SmoM2 mice lead to the rapid development of BCC, with lesions appearing two weeks following tamoxifen primarily in the snout, ears and tail. These lesions are characterized by the expansion of basal keratinocytes expressing cytokeratin 14 (K14) (Fig. 1B) and show nuclear staining for p63 and YAP1 (Fig. 1C).

**Fig. 1:**
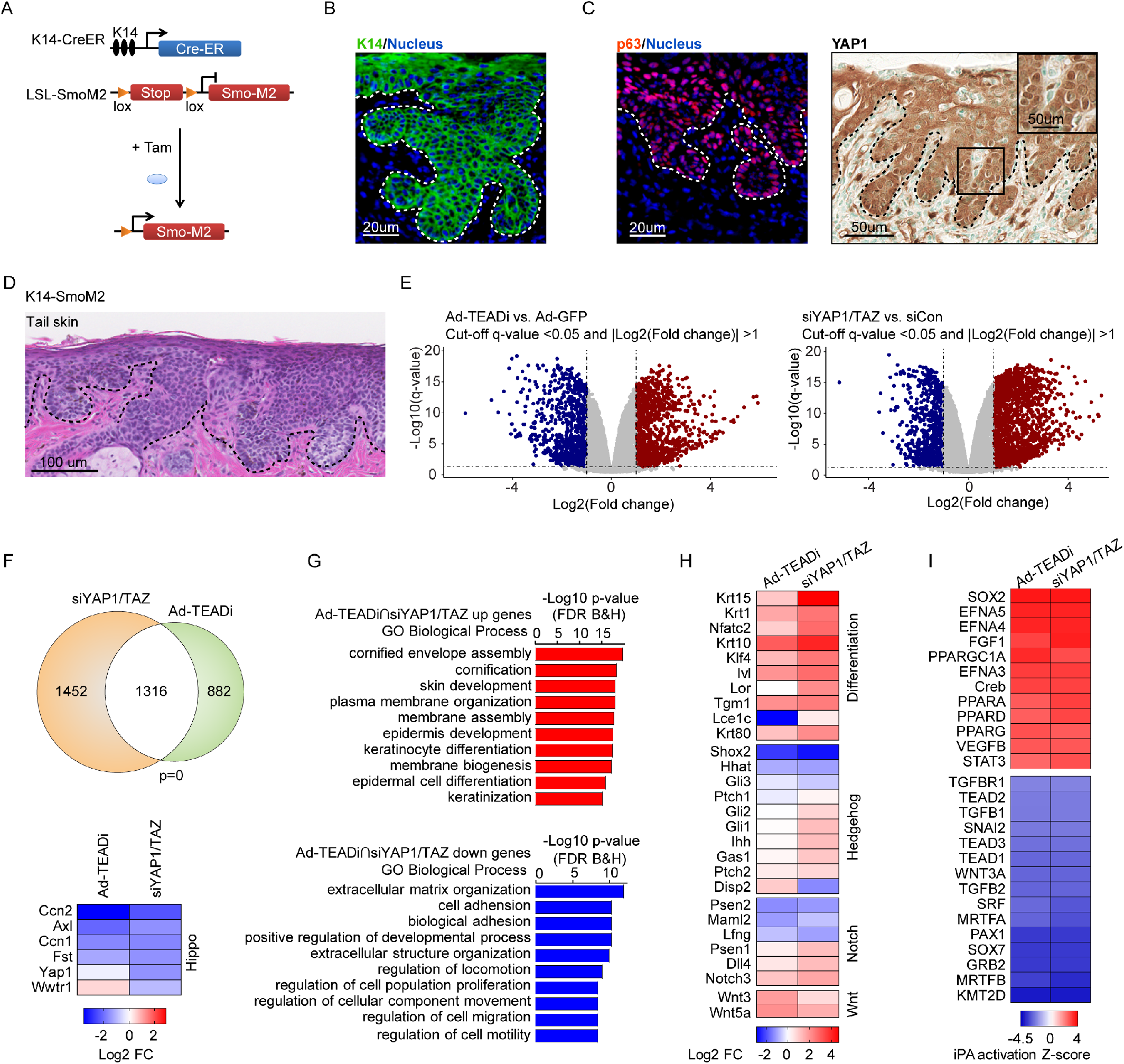
TEAD transcriptional networks regulate differentiation in BCC. **A-** Schematic representation of the K14-SmoM2 tamoxifen (Tam) inducible BCC mouse model. **B and C-** Staining with the indicated markers of K14-SmoM2 BCC tumors 5 weeks after tamoxifen (Tam) induction. Basal layer of epidermis is indicated with dotted line. **D-** H&E staining of tail skin from K14-SmoM2 showing BCC tumors cover the entire epidermis. **E-** Volcano plots from RNAseq from BCC cells treated with TEADi or siYAP1/TAZ, n= 3 for each condition; dotted lines indicate significant genes (q<0.05 and |FC|≥2). **F-** Venn diagram showing the overlap between differently regulated genes from E and heatmap indicating fold change (log2 FC) of YAP1/TAZ and target genes; p-value of the overlap is indicated, Fisher’s exact test. **G-** Top gene Ontology (GO) biological process terms for genes enriched in upregulated (red, q<0.05 and FC≥2) or downregulated (blue, q<0.05 and FC≤-2) genes differentially regulated by both siYAP1/TAZ and TEADi (indicated in white in F). **H-** Heatmap showing fold change of selected genes related to differentiation, Hedgehog, Notch, and Wnt signaling pathways. **I-** Heatmap showing activation Z-score for upstream transcriptional regulators calculated using Ingenuity Pathway Analysis (IPA) for genes in siYAP1/TAZ and TEADi datasets (q<0.05 and |FC|≥2).

While the role of YAP1 and TAZ in the development and progression of BCC lesions in K14-SmoM2 mice has been described before ^5,6^, the precise effects of TEAD inhibition in BCC are not clear. To assess TEAD role in BCC, we utilized the TEAD-inhibitor TEADi and pooled-small interfering RNAs targeting YAP1 and TAZ (siYAP1/TAZ) in isolated skin keratinocytes from the tail of BCC mice 5 weeks following tamoxifen induction. As shown in Figure 1D, at this time point BCC lesions covered the entire tail epidermis. RNA sequencing (RNA-seq) of BCC keratinocytes with adTEADi or siYAP1/TAZ revealed the dysregulated expression of numerous transcripts (Fig. 1E and Table S1). A significant overlap between differentially regulated genes in both conditions was observed (Fig. 1F), indicating common gene networks in TEADi and siYAP1/TAZ datasets. Both TEADi and siYAP1/TAZ led to the significant downregulation of the known TEAD targets *Ccn1* and *Ccn2* (also known as *Cyr61* and *Ctgf* respectively), and *Axl* and *Fst* (Fig. 1F). siYAP1/TAZ lead to a significant reduction on *Yap1* and Taz (*Wwtr1*) expression, while TEADi did not significantly altered these genes (Fig. 1F). Gene ontology (GO) analysis of genes regulated by both TEADi and siYAP1/TAZ indicated an upregulation of processes related to skin and keratinocyte differentiation, while downregulated gene networks were enriched for terms associated with extracellular matrix organization, cell adhesion and migration (Fig. 1G).

Although BCC is characterized by the dysregulation of Hedgehog signaling, we did not find significant alterations on the expression of *Gli* or *Ptch* or other Hedgehog targets by TEADi (Fig. 1H). Multiple alterations were present in genes central for keratinocyte differentiation, including *Klf4, Foxn1, Notch3, Ivl* and *Krt10*, among others (Fig. 1H). A broader analysis of differentiation gene expression revealed that both TEADi and siYAP1/TAZ lead to an activation of differentiation markers related to every differentiation stage (Fig. S1A) and a significant correlation with epidermal cell differentiation and keratinization terms (Fig. 1G). These results indicate that activation of YAP1/TAZ-TEAD transcriptional networks are essential for Hedgehog-mediated blockage of differentiation and tumorigenesis in BCC. TEADi and siYAP1/TAZ downregulated genes showed an enrichment for extracellular matrix and cell adhesion family members (Fig. 1G). Components of the basal membrane, including laminins and collagens were downregulated by siYAP1/TAZ and TEADi, while several metalloproteases were upregulated (Fig. S1B), suggesting that TEAD transcription participates in regulating integrity of the basal membrane and cell adhesion. Adhesion to the basal membrane is closely related to maintenance of cell renewal in keratinocytes and the downregulation of these gene networks could be contributing to the observed differentiation in BCC TEADi cells.

Ingenuity pathway analysis (IPA) of upstream transcriptional regulators affected by siYAP1/TAZ and TEADi confirmed the downregulation of networks related to TEAD transcription factors (Fig. 1I). In addition, genes related to SRF/MRTF were downregulated (Fig. 1I). SRF and MRTF have been linked to therapeutic resistant in BCC ^19^. On the other hand, networks related to ephrins (EFN) and peroxisome proliferator activated receptors (PPAR) were activated (Fig. 1I). TAZ can regulate PPARγ ^20^ and PPAR activation has been implicated in keratinocyte differentiation ^21^. Ephrins have also been involved in reducing keratinocyte proliferation and increased differentiation ^22^. Interestingly, IPA analysis revealed a similarity between the transcriptional networks activated by YAP1/TAZ knockdown and TEADi to those activated by cAMP and PKA signaling (Fig. S1C). This suggests that the transcriptional regulation downstream of PKA is partially mediated by YAP1/TAZ and TEAD blockage.

Overall, our results show that YAP1 and TAZ TEAD-dependent transcriptional networks are involved in the maintenance of the undifferentiated state of BCC cells and that TEAD inhibition leads to the rapid activation of differentiation genes.

### KLF4 transcriptional networks are activated by YAP1/TAZ and TEAD inhibition in BCC

To better understand the dysregulation of differentiation pathways triggered by TEAD blockage in BCC we performed a transcription factor binding site enrichment analysis in proximal promoters of genes regulated by both TEADi and siYAP1/TAZ. Over-represented transcription factor binding sites in downregulated genes included TEAD and SRF (Fig. 2A), supporting a direct role of TEAD in regulating expression of these transcripts. Binding sites in genes upregulated by YAP1/TAZ-TEAD inhibition showed a clear enrichment for KLF4 (Fig. 2A). KLF4 is a central regulator of keratinocyte differentiation ^23,24^ and our group has previously shown that YAP1/TAZ-TEAD regulate the activity of this transcription factor in normal keratinocytes ^15^.

**Fig. 2:**
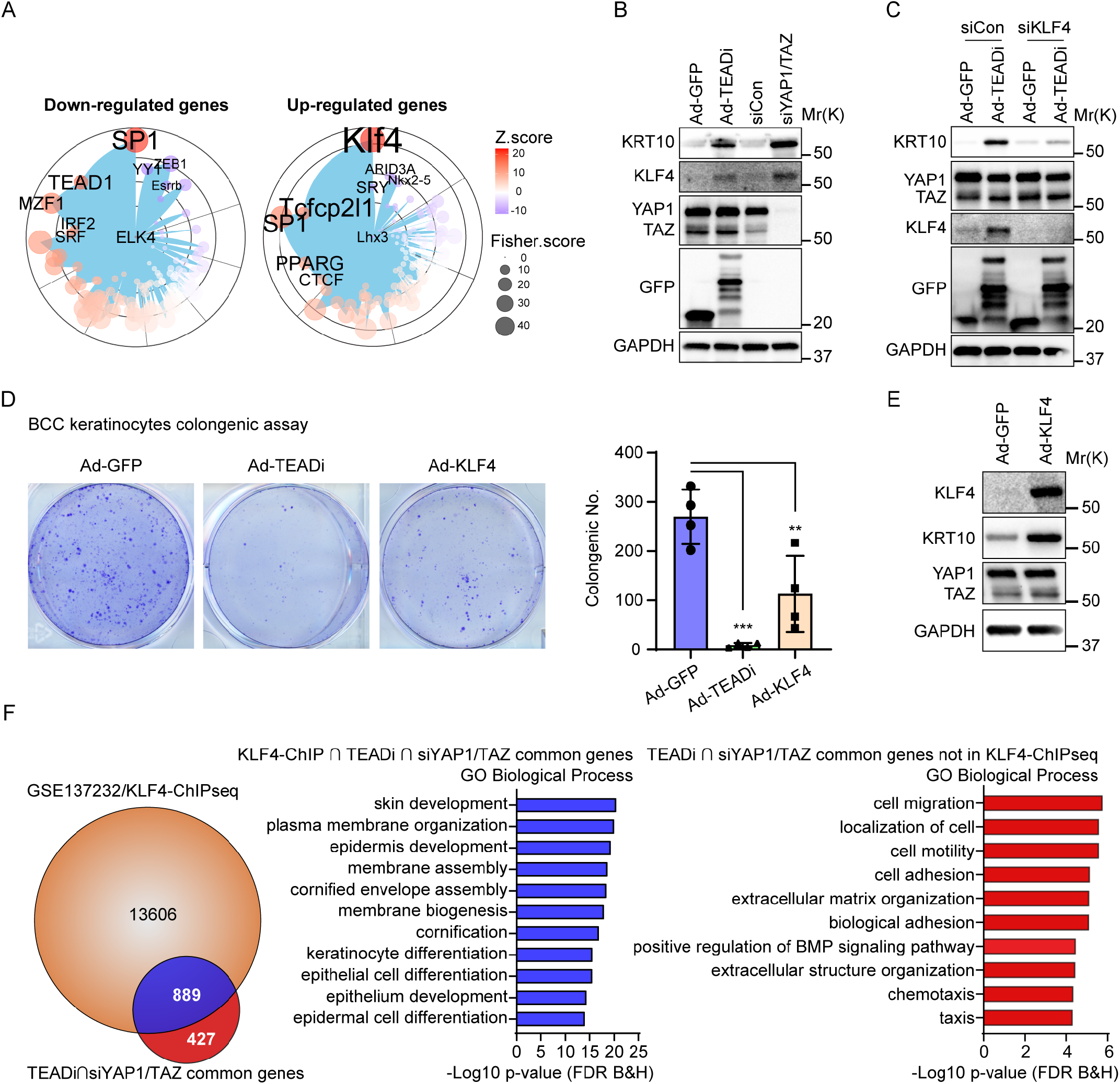
YAP1/TAZ and TEAD inhibition in BCC activate KLF4 transcription. **A-** Graph showing the transcription factor binding site enrichment analysis using genes differentially regulated by both siYAP1/TAZ and TEADi (indicated in white in Fig. 1F); only the top enriched transcription factors are highlighted, color represents Z-score and size represents Fisher-score. **B-** Western blot analysis of the indicated markers 48hs after transduction of cultured BCC keratinocytes with Ad-TEADi and control Ad-GFP, or with pooled siYAP1/TAZ or control siCon. **C-** Western blot analysis of the indicated markers in BCC cultured keratinocytes infected with Ad-TEADi or control Ad-GFP, transduced 12hs later with siRNAs targeting KLF4 or negative control (siCon) and harvested after 48hs. **D-** Clonogenic assay of cultured BCC keratinocytes transduced with control ad-GFP, ad-TEADi or ad-KLF4. Representative picture and colony numbers from 4 replicates. Two-way ANOVA with Dunnett’s multiple comparisons test (**p<0.01, ***p<0.001). **E-** Western blot of the indicated markers in BCC cultured keratinocytes transduced with Ad-KLF4 or control ad-GFP for 48hs. **F-** Venn diagram indicating the overlap between genes with KLF4 binding (KLF4 ChIP-seq) and genes commonly regulated by TEADi and siYAP1/TAZ; and top GO biological process terms from genes in the overlap area (indicated in blue) or exclusively in the TEADi-siYAP1/TAZ dataset (indicated in red).

BCC keratinocytes transduced with TEADi or siYAP1/TAZ showed an upregulation of KLF4 expression and the differentiation marker KRT10 (Fig. 2B), which could be prevented by downregulation of KLF4 by pooled-small interfering RNAs (siKLF4, Fig. 2C). Furthermore, BCC keratinocytes transduced with adenoviruses expressing TEADi (Ad-TEADi) or KLF4 (Ad-KLF4) presented reduced clonogenic capacity compared with keratinocytes transduced with control GFP (Ad-GFP, Fig. 2D), and KLF4 overexpression was sufficient to induce the expression of KRT10 in these cells (Fig. 2E). These results validate the necessity of TEAD activation and KLF4 inhibition for cell growth in BCC cells.

In order to validate the core function of KLF4 in BCC keratinocyte differentiation, we utilized available information on KLF4 chromatin immunoprecipitation analysis from mouse epidermis (ChIP-seq) ^25^. Numerous genes differentially regulated by both TEADi and siYAP1/TAZ showed KLF4 binding (Fig. 2F). GO enrichment analysis demonstrated that these KLF4 bound genes were related to skin development and keratinocytes differentiation, while the set of genes without KLF4 binding did not show enrichment for differentiation terms (Fig. 2F). Our results indicate that repression of KLF4 transcriptional networks by YAP1/TAZ-TEAD is essential for maintaining basal cell identity and block differentiation in BCC.

### YAP1 and TAZ regulate inflammatory related gene networks independently of TEAD

We next focused on the TEAD independent functions of YAP1/TAZ in BCC by evaluating pathways preferentially regulated by siYAP1/TAZ but not TEADi. Interestingly, YAP1 and TAZ knockdown lead to a significant upregulation of *Gli1*, the Hedgehog regulating gene *Gas1* and Indian hedgehog (*Ihh*) (Fig. 1H), indicating the possibility of a TEAD-independent crosstalk between Hedgehog and Hippo signaling. siYAP1/TAZ also lead to an increase in *Wnt5a* and *Wnt3*, which could suggest an upregulation of the Wnt pathway, although global changes in this pathway were not clear (Fig. 1H). IPA analysis for YAP1/TAZ and TEADi regulated genes indicated preferential activation by YAP1/TAZ knockdown of several networks related to NF-κB signaling (Fig. 3A). YAP1 has been described to reduce NF-κB activation and the concomitant expression of proinflammatory cytokines IL-6, TNF-α and IL-1β ^26^, and our analysis showed that siYAP1/TAZ but not TEADi increases IL-6, TNF and IL-1 gene networks (Fig. 3A). Other immune-modulatory pathways were also preferentially activated by YAP1 and TAZ knockdown, including interferon γ (IFNγ) and STAT3 signaling (Fig. 3A). Transduction of BCC keratinocytes with siYAP1/TAZ and TEADi confirmed the differential activation of Stat3 (Fig. 3B).

**Fig. 3.**
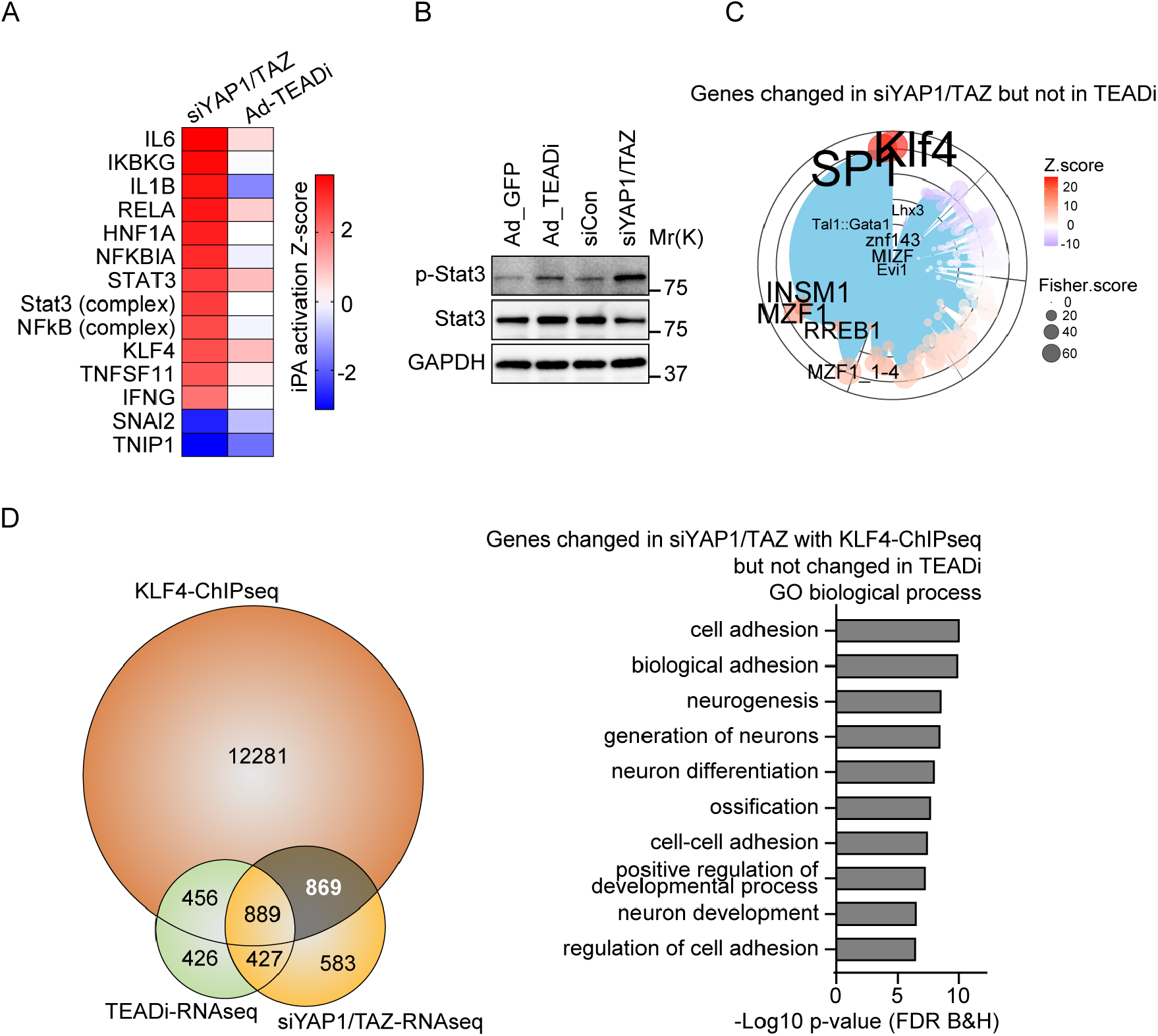
YAP1 and TAZ regulate inflammatory signals via TEAD independent gene networks. **A-** Heatmap showing activation Z-score for selected IPA upstream transcriptional regulators for genes differentially regulated in siYAP1/TAZ or TEADi datasets (q<0.05 and |FC|≥2). **B-** Western blot showing the expression level of phosphorylated (p-) and total Stat3 in BCC cells treated with the indicated conditions. **C-** Graph showing the transcription factor binding site enrichment analysis in genes exclusively regulated by siYAP1/TAZ (indicated in gray in D); only the top enriched transcription factors are highlighted, color represents Z-score and size represents Fisher-score. **D-** Venn diagram indicating the overlap between KLF4 ChIP-seq target genes and genes differentially regulated by siYAP1/TAZ and TEADi; and selected GO biological process terms from area highlighted in gray.

Interestingly, IPA analysis also indicated a stronger activation of KLF4 gene networks in the siYAP1/TAZ knockdown dataset (Fig. 3A). Analysis of over-represented transcription factor binding sites in genes differentially regulated by siYAP1/TAZ but not TEADi also showed an enrichment for KLF4 (Fig. 3C). It is worth noting that this TEADi independent set of genes did not present enrichment for TEAD binding sites (Fig. 3C). Cross-reference of the TEADi and siYAP1/TAZ differential regulated genes with KLF4-ChIPseq data indicated that KLF4 networks were activated in both conditions (Fig. 3D). Genes exclusively regulated by YAP1/TAZ that present KLF4 binding were not related to epithelial differentiation but to cell adhesion (Fig. 3D). These results suggest a TEAD dependency for the activation of KLF4 differentiation networks resulting from YAP1/TAZ inhibition.

### TEADi leads to rapid elimination of tumor cells in BCC lesions

To study the effect of TEADi in BCC tumors we crossed K14-SmoM2 mice with mice carrying LSL-rtTA ^27^ and tetracycline-inducible TEADi (TRE-TEADi) ^15^ (Fig. 4A). The resulting animal model allowed us to trigger BCC formation with tamoxifen while controlling the expression of TEADi by feeding mice doxycycline chow at different points during tumor development (Fig. 4A).

**Fig. 4:**
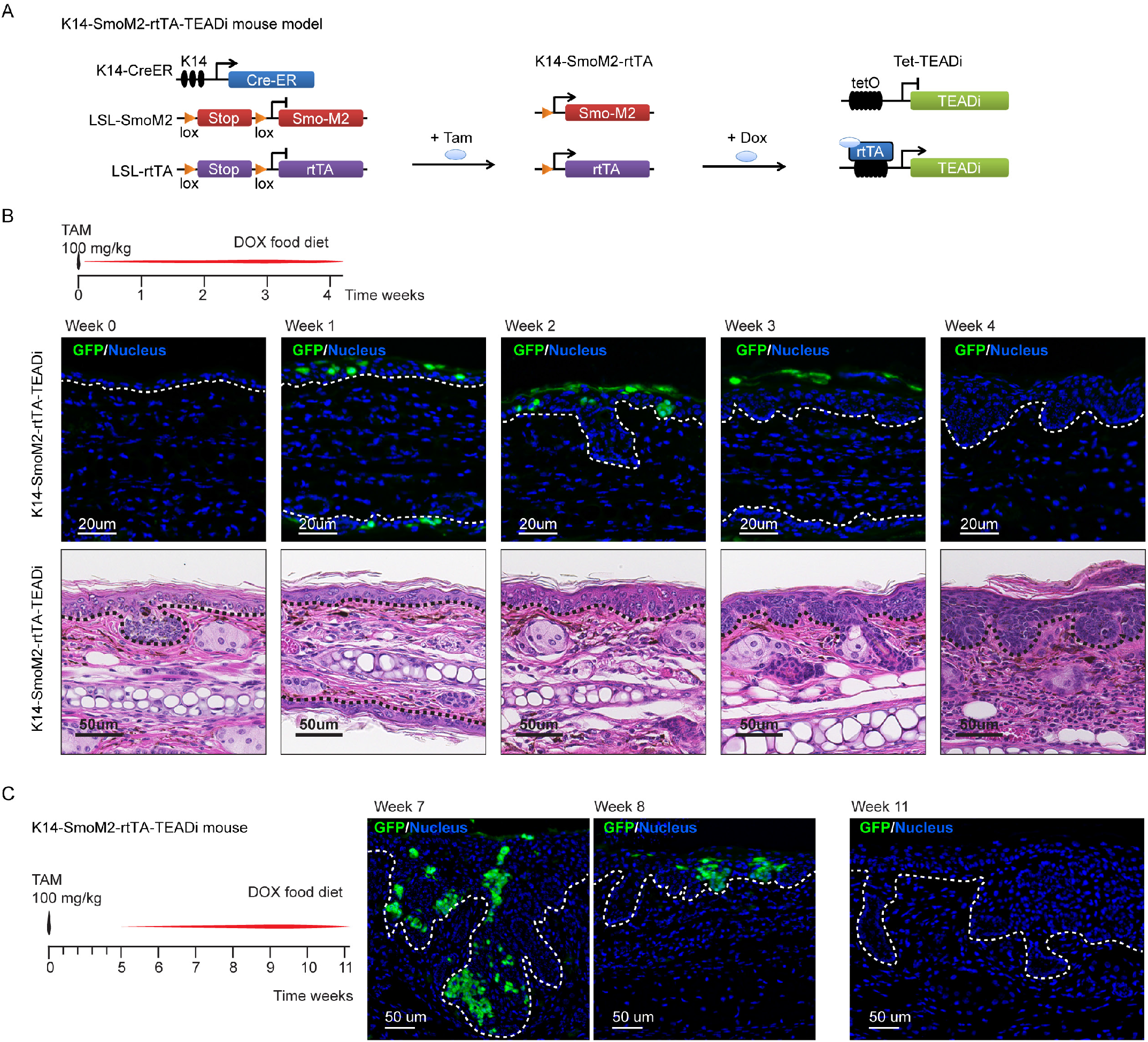
TEAD inhibition leads to rapid elimination of tumor cells in BCC lesions. **A-** Schematic diagram presenting the mouse model used to trigger TEADi expression in BCC tumors. **B-** Timeline and histological analysis of mice induced with one dose of tamoxifen and concomitantly given doxycycline to activate TEADi expression; GFP staining shows TEADi positive cells at the indicated time points; representative pictures from mice analyzed: n=3 for week 0, 1 and 3; n=4 for week 2 and 4. **C-** Timeline and TEADi expression (GFP) in mice induced with one dose of tamoxifen and given doxycycline 5 weeks later to activate TEADi expression following tumor formation; representative pictures from n=4 mice analyzed for weeks 7, 8 and 11. Basal layer of epidermis is indicated with dotted line. Tam: tamoxifen, Dox: doxycycline food.

When mice were induced with a single dose of tamoxifen and concomitantly fed doxycycline chow, we observed scattered cells in the epidermis positive for TEADi (GFP, Fig. 4B). Over time, TEADi expressing cells were eliminated from the epidermis and localized to the upper and differentiated layers of the skin by week 3 (Fig. 4C). Although BCC lesions still arise in these mice, the resulting tumors were smaller and TEADi negative (Fig. 4C and S2), indicating that cells in which TEAD was blocked did not proceed to form tumors. The appearance of BCC lesions negative for GFP in this model could be due to incomplete recombination of LSL-rtTA in a subset of cells that express SmoM2, allowing these cells to proceed with tumor formation. A similar scenario was reported in BCC studies with YAP1/TAZ knockout animals ^5,6^, in which BCC tumors arise in YAP1/TAZ and YAP1 knockout mice that are positive for YAP1 due to incomplete recombination.

In another set of experiments, mice were treated with tamoxifen and TEADi expression was induced 5 weeks later. Although in this case BCC lesions were already developed before expression of the inhibitor, we observed a similar trend in which all TEADi positive cells were eliminated from BCC tumors over time (Fig. 4D). Analysis of differentiation and proliferation markers indicated that intramural TEADi cells were positive for the differentiation markers keratin 10 (KRT10) and KLF4 (Fig. 5A and B) and showed reduced labeling for the proliferation marker PCNA (Fig. 5C).

**Fig. 5:**
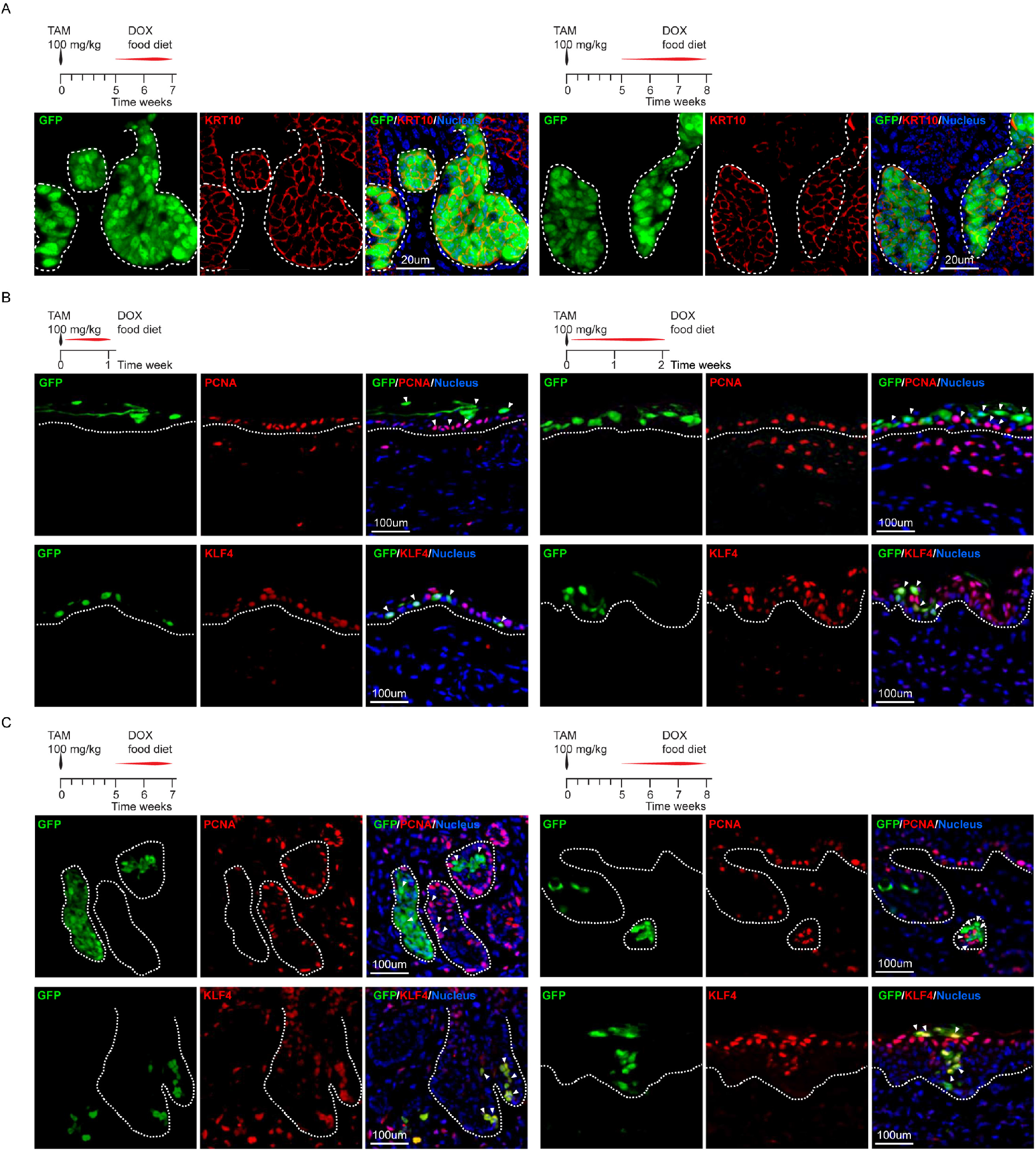
TEAD inhibition induces differentiation and reduces proliferation in BCC cells. **A to C-** Immunofluorescence analysis of the differentiation markers KRT10 and KLF4, the proliferation marker PCNA and TEADi (GFP). Timeline indicates treatment regimen and time-point when tissue was harvested. Representative pictures from three mice analyzed. Basal layer of epidermis is indicated with dotted line. TAM: tamoxifen, DOX: doxycycline food.

Our in vivo data shows that inhibition of TEAD during or subsequent to BCC tumor formation leads to the rapid activation of differentiation pathways and elimination of cells from tumor lesions and validate our conclusion that activation of TEAD is necessary for preventing differentiation downstream from Hedgehog signaling in BCC.

## Discussion

In this study we use knockdown strategies and specific transcriptional inhibition to discern the TEAD-dependent and independent effects of YAP1/TAZ in cancer. We find that blockage of TEAD leads to rapid differentiation of BCC cells by activating transcriptional networks centered on KLF4, demonstrating a TEAD dependency for the blockage of differentiation downstream of Smo in BCC. While YAP1/TAZ knockdown recapitulates TEAD inhibition, it results in additional effects in cancer cells. siYAP1/TAZ but not TEADi leads to the activation of INFγ, STAT and NF-κB gene networks as well as increases in the levels of some Hedgehog pathway members. Reduced INF and STAT signaling has been attributed to cytoplasmic effects mediated by YAP1 and TAZ ^28,29^. YAP1 has also been shown to dampen NF-kB activation by interacting directly with TRAF6 ^26^ and TAK1 ^30^. Together, these data indicate that TEAD-independent effects of YAP1 and TAZ might be mediated mainly through cytoplasmic rather than nuclear interactions with other proteins.

TEAD inhibition in BCC leads to differentiation and elimination of tumor cells, making TEAD inhibitors candidates for differentiation therapy ^31^. Nevertheless, our data indicates that cells that evade the activation of differentiation pathways could be less sensitive to TEAD blockage. Furthermore, potential tumor promoting and immune evasion signals mediated by cytoplasmic interactions of YAP1 and TAZ could pose a challenge on targeting this pathway, since blocking TEAD would not be able to prevent non-nuclear effects of YAP1/TAZ. Inhibitors that target cytoplasmic and nuclear functions of YAP1 and TAZ could prove to be more effective for cancer treatment. In this regard, we found that transcriptional networks activated by YAP1/TAZ knockdown present high similarity with those activated by cAMP and PKA signaling. Indeed, cAMP and PKA blockage can lead to BCC formation by inducing the cell-autonomous activation of GLI and YAP1 ^32^. Activation of PKA on the other hand can lead to blockage of YAP1 function ^32-34^. Forskolin, which results in increased intracellular cAMP levels and PKA activation, has been shown to reduce BCC tumor formation in mouse models ^35^, highlighting the potential of this pathway for YAP1/TAZ inhibition and BCC treatment.

We also find that KLF4 is central for the activation of differentiation gene networks following TEAD inhibition. Although KLF4 genomic alterations are not common in skin cancer, KLF4 is downregulated in BCC and SCC tumors and KLF4 knockout mice present increase sensitivity to tumor formation ^23^. KLF4 is also responsible for reduced self-renewal and increased differentiation of skin cancer-initiating cells in mouse models ^36^. Since KLF4 expression can also result in the concomitant inhibition of TEAD ^15^, finding avenues to regain or increase KLF4 activity in BCC could prove effective in therapeutic strategies for these tumors.

The activation of KLF4 and differentiation pathways resulting from TEAD inhibition in BCC cells closely recapitulates our findings in normal human keratinocytes and mouse skin ^15^, rising potential limitations with this tumor model. While oncogene-driven models like SmoM2 provide invaluable information on cancer initiation and progression, they do not completely reflect the complex mutational landscape of tumors. Particularly relevant is the fact that carcinomas accumulate numerous genomic and epigenomic modifications that render them incapable of differentiation, compared with single oncogene models that do not acquire the necessary mutations and promoter hypermethylation to completely abrogate differentiation. While our data suggests that TEAD inhibition in normal and mutant Smo cancer cells would trigger their elimination by activating differentiation, more research is needed to understand the networks resulting from TEAD inhibition in more transformed cells.

## Methods

### Cell culture, transfections and adenoviral transductions

Cells were cultured at 37°C in the presence of 5% CO2. BCC tumor cells were isolated from tail skin of K14-SmoM2 mice five weeks following tamoxifen induction. Skin was disinfected with 10% iodine in phosphate saline buffer (PBS) and incubated with 2U/ml of dispase (STEMCELL technologies) overnight at 4°C. Next day, epidermis was scraped with forceps and digested with trypsin 0.05%+EDTA solution (SIGMA) at 37°C for 4min. Cells were filtered using a 100µm cell strainer, pelleted at 400g and plated in collagen I coated dishes (0.3 mg/ml collagen I, Corning, in 1% acetic acid). BCC cells were cultured using EpiLife medium with 60 µM calcium (GIBCO MEPI500CA), supplemented with defined growth supplement (EDGS, GIBCO S0125), mouse EGF (10ng/ml, R&D Systems 2028EG200), and Y-27632 compound (10 µM, Tocris Bioscience 12-541-0). For siRNA experiments, cells were transfected with the corresponding siRNAs one day after plating and treated/harvested 48hs after transfection. siRNAs were siGENOME SMARTpool from Dharmacon/Horizon, siYAP1 (M-046247-01-0010), siTAZ (*Wwtr1*, M-041057-01-0010), siKLF4 (Klf4, M-040001-02-0010) and non-targeting control siRNA (D-001206-13). siRNA was transfected at a concentration of 8 pmol cm-2 using Lipofectamine RNAiMAX (Invitrogen) according to manufacturer’s instructions. For adenoviral transduction, cells were incubated with a multiplicity of infection (MOI) of 25 with adeno-GFP (control) or adeno-TEADi for the indicated times. TEADi adenoviruses were produced, purified and titered by Vector Biolabs in an adenoviral-Type 5 (dE1/E3) backbone with a CMV promoter. GFP control and KLF4 adenoviruses were purchased from Vector Biolabs (Ad-CMV-GFP, catalog no. 1060; Ad-CMV-KLF4, catalog no. 1787). To assess colony-forming efficiency, equal number of keratinocytes from corresponding mice were infected with adeno-GFP or adeno-TEADi and plated in triplicate in six-well plates and grown for 5 to 7 days. Plates were fixed using 4% paraformaldehyde for 15 mins and stained with crystal violet (0.5%, SIGMA).

### Gene Expression Analysis

RNA was isolated using TRIzol Reagent (Invitrogen) according to manufacturer’s instruction. Cells were lysed using Precellys lysing kit (Bertin Instruments) and mRNA integrity was validated with Agilent TapeStation system. 3 independent samples were sequenced for each condition. mRNA expression profiling was performed in the CCR-Sequencing Facility at the NIH. Reads of the samples were trimmed for adapters and low-quality bases using Trimmomatic software before alignment with the reference genome mouse-mm10 and annotated transcripts using STAR. Gene counts were filtered by genes with ≥5 reads and normalized to TMM (Trimmed Mean of M values) using Partek Flow software, version 8 (Partek Inc). TMM normalized counts were used for differential analysis using PARTEK Flow GSA algorithm (Partek Inc). Gene Ontology (GO) terms were obtained with ToppGene ^37^ using indicated gene sets. Canonical pathways and upstream regulators analysis were generated with Ingenuity Pathway Analysis (IPA, Ingenuity Systems, www.ingenuity.com) by using genes with q<0.05 and |FC|≥2.0. Analysis of over-represented conserved transcription factor binding sites was performed with oPOSSUM ^38^ using upregulated (q<0.05, FC≥2.0) and downregulated (q<0.05, FC≤-2.0) genes, looking at 2kb upstream/downstream sequence with a conservation cutoff of 0.6. Differentiation gene clusters and functional signature genes for cell adhesion and ECM were obtained from ^39^.

### Mice

All animal studies were carried out according to approved protocols from the NIH-Intramural Animal Care and Use Committee (ACUC) of the National Cancer Institute, in compliance with the Guide for the Care and Use of Laboratory Animals. TRE-TEADi mice were described before ^15^. Other mouse lines were obtained from The Jackson Laboratory: LSL-SmoM2 (stock 005130), LSL-rtTA (stock 005670) and K14CreERT (stock 005107). Both LSL-SmoM2 and LSL-rtTA transgenes are in the same Rosa26 locus. To generate K14CreERT^+/-^ LSLSmoM2^+/-^ LSL-rtTA^+/-^ TRE-TEADi^+/-^ mice, homozygous K14-SmoM2 mice were crossed with LSL-rtTA homozygous-TRE-TEADi heterozygous mice. Both male and female mice were used, and all experiments were conducted using littermate controls. Housing conditions were as follow: temperature set point is 72±4 °F (22.2±2.2 °C), light cycle of 12hs on 6am to 6pm and 12hs off, NIH-03I rodent diet. Tamoxifen (SIGMA) was dissolved in ethanol and mixed in oil (Miglyol 810N, Peter Cremer North America LP) for intraperitoneal injection in mice at a dose of 100mg/kg. Doxycycline was administered in food grain-based pellets (Bio-Serv) at 6g/kg.

### Immunofluorescence and Immunohistochemistry

Immunofluorescence analysis of mouse skin was performed on tissue sections embedded in paraffin. Sections were rehydrated and prepared for staining by antigen retrieval in 10mM Sodium Citrate buffer pH 6, washed and blocked with 3% BSA for 1hr at room temperature. Slides were incubated with the primary antibody overnight at 4°C, washed three times with PBS followed by incubation with the secondary antibody for 1.5hs at room temperature. Sections were mounted in FluorSave Reagent (Millipore, #345789) with #1.5 coverslips for imaging. Nuclei were stained with Hoechst 33342 (1:2000, Invitrogen, #H3570). The following antibodies were used: GFP (Aves Labs; catalogue no. GFP-1010; 1:500), KRT10 (BioLegend; catalogue no. 905401; 1:400), PCNA (Cell Signaling; catalogue no. 13110S; 1:400), p63 (Cell Signaling; catalog no. 39692S; 1:400), mouse KLF4 (R&D; catalogue no. AF3158; 1:200), KRT14 (BioLegend; catalogue no. 906001; 1:400). The secondary antibodies were Donkey anti-Rabbit IgG Alexa Fluor 555 (Invitrogen; catalog no. A-31572; 1:1000), Donkey anti-Goat IgG Alexa Fluor 546 (Invitrogen; catalog no. A-11056; 1:1000), Goat anti-Chicken IgY Alexa Fluor 488 (Invitrogen; catalog no. A-11039; 1:1000). Images were obtained using a Keyence BZ-X700 with automatic stage and focus with BZX software (objective CFI Plan Apo λ20x NA 0.75, Nikon). Final images were bright contrast adjusted with BZX analysis software (Keyence) or PowerPoint. For histological analysis, tissues were embedded in paraffin and 3-μm sections were stained with H&E. Stained H&E slides were scanned at 40x using an Aperio CS Scanscope (Aperio). Tumor burden in ear skin was quantified in H&E sections with Image J (imagej.nih.gov) by calculating the tumor fraction versus total epidermis in regions of 256×256 pixel squares. Immunohistochemistry staining of mouse skin was performed on tissue sections embedded in paraffin. Sections were prepared for staining by antigen retrieval in 10mM Sodium Citrate buffer pH 6, washed and blocked with 3% BSA for 1hr at room temperature. Slides were then incubated with the primary antibody YAP1 (Cell Signaling; clone no. D8H1X; catalogue no. 14074; 1:100) overnight at 4°C, washed three times with PBS. Secondary antibody incubation and HRP reaction was conducted with Vectastain ABC kit HRP (Vector Laboratories).

### Immunoblot Analysis

For Western blot cells were lysed by sonication at 4°C in lysis buffer (50 mM Tris-HCl, 150 mM NaCl, 1 mM EDTA, 1% Nonidet P-40, 0.5% Sodium Deoxycholate, 0.1% SDS) supplemented with complete protease inhibitor cocktail (Roche, #6538304001) and phosphatase inhibitors (PhosSTOP, Sigma-Aldrich, #4906837001). Equal amounts of total cell lysate proteins were subjected to SDS-polyacrylamide gel electrophoresis and transferred to PVDF membranes. Primary antibodies used were: anti-GAPDH (Cell Signaling; clone no. 14C10; catalogue no. 2118; 1:2000), anti-GFP (Cell Signaling; clone no. D5.1; catalogue no. 2956; 1:2000), KRT10 (BioLegend; catalogue no. 905401; 1:1000), Stat3 (Cell Signaling; clone no. D1B2J; catalogue no. 30835; 1:1000), p-Stat3 (Tyr705) (Cell Signaling; clone no. D3A7; catalogue no. 9145; 1:1000), KLF4 (R&D; catalogue no. AF3158; 1:1000), YAP1 (Cell Signaling; clone no. D8H1X; catalogue no. 14074; 1:1000), and TAZ (Cell Signaling; clone no. V386; catalogue no. 4883; 1:1000). Secondary HRP-conjugated antibodies used were: Pierce peroxidase goat antimouse IGG (H+L) (ThermoFisher, catalogue no. 31432; 1:4000) and Pierce peroxidase goat antirabbit IGG (H+L) (ThermoFisher, catalogue no. 31462; 1:4000). Secondary antibody was incubated at RT for 1 hr. Bands were detected using a ChemiDoc Imaging System (Bio-Rad) with Clarity Western ECL Blotting Substrates (Bio-Rad) according to manufacturer’s instructions. Blot images were processed using ImageLab software v5.2.1 (Bio-Rad).

### Statistics

All analyses were performed in triplicate or greater and the means obtained were used for ANOVA or independent t-tests. Statistical analyses were carried out using the Prism 7 statistical analysis program (GraphPad). Statistical analysis of intersections in Venn diagrams was performed by hypergeometric test (one tailed Fisher’s exact test). Asterisks denote statistical significance.

### Data availability

RNAseq primary and processed data generated in this manuscript is available from GEO database GSE156913 (BCC data). Processed RNAseq data is provided in Supplementary Tables S1. RNAseq data from normal keratinocytes is from GSE136876 ^15^. KLF4 ChIP-seq data is from ^25^.

## Acknowledgments

This research was supported by the Intramural Research Program of the National Institutes of Health, National Cancer Institute, Center for Cancer Research (ZIA BC 011764 and ZIA BC 011763). This work used the computational resources of the NIH High-Performance Computing Biowulf Cluster. We thank Jennifer E Dwyer for assistance with slide scanning in the Aperio CS Scanscope, and members of the CCR Sequencing Facility at Frederick National Laboratory for Cancer Research for their help during sample preparation, sequencing and data processing.

## Author contributions

R.I.B. initiated the study; Y.Y. and R.I.B. designed the study and experiments; Y.Y., N.S.P and R.I.B. performed experiments; Y.Y. and R.I.B. analyzed and interpreted data; R.I.B. and Q.C. provided administrative, technical, and material support; Y.Y. and R.I.B. prepared the manuscript.

## Additional information Competing interests

The authors declare no competing interests.

