## Supplementary Figures for "YAP1-TAZ/TEAD transcriptional networks restrain differentiation downstream of oncogenic Hedgehog-SMO activity"

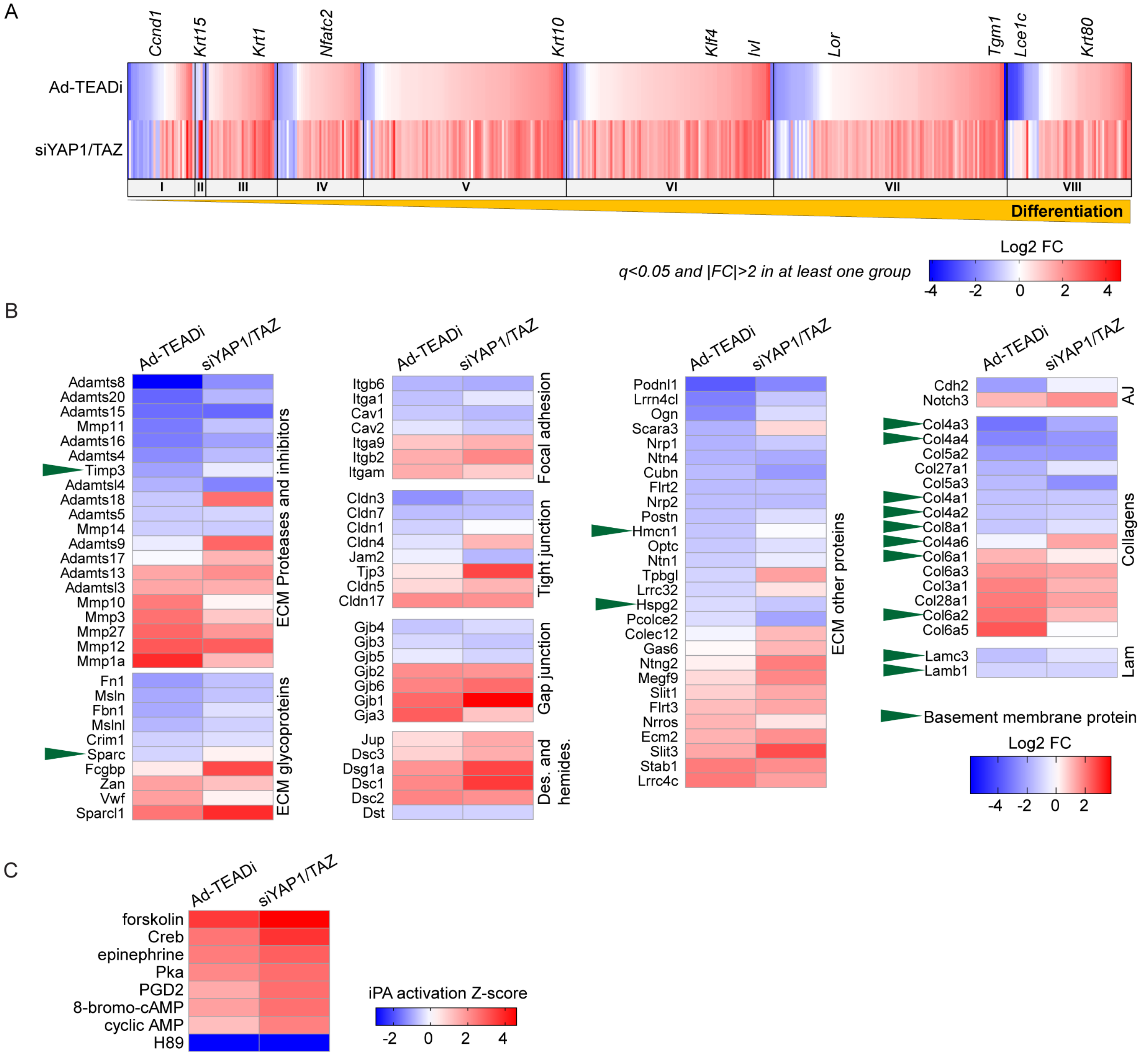


**Fig. S1. Effects of TEAD inhibition in BCC cells. A-** Heatmap showing fold change of differentiation markers in TEADi and siYAP1/TAZ datasets (q<0.05 and |FC|≥2 in at least one condition); clusters of differentiation genes from basal (I) to terminal differentiation (VIII) and key differentiation markers are indicated. **B-** Expression changes in extracellular matrix and cell adhesion components in TEADi and siYAP1/TAZ datasets (q<0.05 and |FC|≥2 in at least one condition). Basement membrane proteins are highlighted by arrows. **C-** Heatmap showing activation Z-score for selected IPA upstream transcriptional regulators related to cAMP and PKA signaling in genes differentially regulated in siYAP1/TAZ or TEADi datasets (q<0.05 and |FC|≥2). Forskolin, epinephrine, and prostaglandin D2 (PGD2) are activators of cAMP and PKA. H89 is an inhibitor of the kinase activity of PKA.

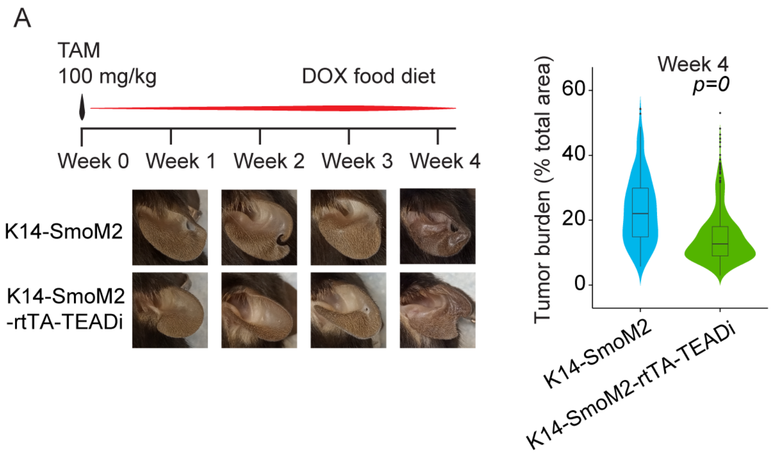


**Fig. S2. TEADi reduces tumor burden in BCC.** Timeline, representative pictures and quantification of ear epidermis tumor burden in mice induced to form BCC tumors concomitantly expressing or not TEADi. For tumor burden K14-SmoM2 n=253 and K14-SmoM2-rtTA-TEADi n=450 individual areas from 3 mice, two-tailed unpaired t test. Ear pictures are from different mice harvested at indicated time-point.

### Supplementary Table

**Table S1**: Differential expression analysis using PARTEK Flow GSA algorithm in BCC cells transduced with GFP (Ad-GFP, control), TEADi (Ad-TEADi), siRNA control (siCon) or siRNA targeting YAP1 and TAZ (siYT) for 48hs. For each condition triplicate samples were used.
